# Genetic and environmental determinants of stressful life events and their overlap with depression and neuroticism

**DOI:** 10.1101/211896

**Authors:** Toni-Kim Clarke, Yanni Zeng, Lauren Navrady, Charley Xia, Chris Haley, Archie Campbell, Pau Navarro, Carmen Amador, Mark J. Adams, David M. Howard, Aleix Arnau Soler, Caroline Hayward, Pippa A. Thomson, Blair H. Smith, Sandosh Padmanabhan, Lynne J. Hocking, Lynsey S. Hall, David J. Porteous, Ian J. Deary, Andrew M. McIntosh

## Abstract

**Background:** Stressful life events (SLEs) and neuroticism are risk factors for major depressive disorder (MDD). However, SLEs and neuroticism are heritable traits that are correlated with genetic risk for MDD. In the current study, we sought to investigate the genetic and environmental contributions to SLEs in a large family-based sample, and quantify any genetic overlap with MDD and neuroticism.

**Methods:** A subset of Generation Scotland: the Scottish Family Health Study, consisting of 9618 individuals comprise the present study. We estimated the heritability of SLEs using pedigree-based and molecular genetic data. The environment was assessed by modelling familial, couple and sibling components. Using polygenic risk scores (PRS) and LD score regression we analysed the genetic overlap between MDD, neuroticism and SLEs.

**Results:** Past 6-month life events were positively correlated with lifetime MDD status (β=0.21, r^2^=1.1%, p=2.5 × 10^−25^) and neuroticism (β =0.13, r^2^=1.9%, p=1.04 × 10^−37^). Common SNPs explained 8% of the variance in personal life events (those directly affecting the individual) (S.E.=0.03, p=9 × 10^−4^). A significant effect of couple environment accounted for 13% (S.E.=0.03, p=0.016) of variation in SLEs. PRS analyses found that individuals with higher PRS for MDD reported more SLEs (β =0.05, r^2^=0.3%, p=3 × 10^−5^). LD score regression demonstrated genetic correlations between MDD and both SLEs (r_G_=0.33, S.E.=0.08) and neuroticism (r_G_=0.15, S.E.=0.07).

**Conclusions:** These findings suggest that SLEs are partially heritable and this heritability is shared with risk for MDD and neuroticism. Further work should determine the causal direction and source of these associations.

## Introduction

The importance of stressful life events (SLEs) in the aetiology of Major Depressive Disorder (MDD) is widely recognised (Kendler, Karkowski, & Prescott, 1999; Kessler, 1997; Surtees et al., 1986). A longitudinal study showed that the odds ratio for the onset of MDD in the month of reporting a SLE is 5.64 (Kendler et al., 1999). Understanding the precise relationship between reporting SLEs and lifetime MDD has, however, proven challenging as factors such as genetics and early environment influence both traits (Kendler & Gardner, 2010).

Whilst SLEs are sometimes considered to be random environmental effects, several studies have shown that reporting SLEs is heritable with estimates from twin studies ranging from 20 to 50% (Billig, Hershberger, Iacono, & McGue, 1996; Foley, Neale, & Kendler, 1996; Kendler, Neale, Kessler, Heath, & Eaves, 1993; Plomin, Lichtenstein, Pedersen, McClearn, & Nesselroade, 1990). SLEs are categorized into dependent events and independent events. Dependent SLEs, such as a relationship problems or job loss, may be, in part, the result of a person’s own behaviour and directly affect the individual. Independent SLEs events, including death or illness of a relative, are more likely to be beyond the control of the individual. The estimated heritability of dependent life events (28-45%) is higher than independent life events (7%) which tend to be more strongly influenced by familial environment (Bemmels, Burt, Legrand, Iacono, & McGue, 2008; Boardman, Alexander, & Stallings, 2011).

Personality can influence the reporting and experience of SLEs. Higher neuroticism levels/scores not only increase risk for MDD but can also moderate the relationship between SLE’s and MDD. A study of 7500 twins found that the depressive effects of SLEs were more pronounced in individuals with higher neuroticism (Kendler, Kuhn, & Prescott, 2004). A four-year longitudinal study of young adults also found that higher neuroticism levels/scores is associated with greater reporting of negative life events (Magnus, Diener, Fujita, & Pavot, 1993).

Greater genetic risk factor for MDD increases the propensity to report SLEs. Twin studies have shown that the risk for SLEs is greater in monozygotic twins with a depressed co-twin compared to dizygotic twins (Kendler & Karkowski-Shuman, 1997). Individuals at genetic risk for MDD may select themselves into high risk environments or have a greater vulnerability to the depressive effects of SLEs (Kendler et al., 1995). This is supported by the observation that depressed individuals tend to experience more dependent SLEs (Chun, Cronkite, & Moos, 2004; Hammen, 1991; Harkness & Luther, 2001; Harkness, Monroe, Simons, & Thase, 1999). As neuroticism is highly correlated with depression, both phenotypically (Jylha & Isometsa, 2006) and genetically (Glahn et al., 2012; Jardine, Martin, & Henderson, 1984), it is also possible that personality traits associated with MDD increase the sensitivity to and/or the reporting of SLEs amongst depressed individuals.

Recent studies have aimed to find the proportion of SLE heritability attributed to common genetic variation using genome-wide SNP data. One study estimated the SNP heritability of SLEs to be 29% (p=0.03, S.E.=0.16) in a sample of 2578 unrelated individuals enriched for MDD cases (Power et al., 2013). However, another study of 7179 African American women found the SNP heritability of SLEs to be only 8% (p=0.02, S.E.=0.04) (Dunn et al., 2016). A significant genetic correlation between SLEs and MDD in African American women was observed by Dunn et al. (r=0.95, p=0.01)(Dunn et al., 2016) using bivariate GCTA-GREML, suggesting that genetic variants that influence MDD risk are also relevant for SLEs. The difference in heritability estimates for SLEs may be the result of differential genetic architectures and familial and environmental effects across samples. Previous studies have shown that more accurate estimates of heritability can be obtained when simultaneously modelling SNP genetic effects in the presence of familial environment (Xia et al., 2016; Zeng et al., 2016). If the correlation between SLEs and MDD can be explained by genetic or familial environmental factors, then this may signpost the most effective strategies for future research by highlighting the optimal periods and opportunities for intervention.

In the present study we aim to estimate the SNP and pedigree heritability of SLEs and also the contribution of couple, sibling and nuclear family effects on SLEs in a family-based cohort drawn from the population of Scotland, Generation Scotland: the Scottish Family Heath Study (GS) (Nagy et al., 2017; Smith et al., 2013; Smith et al., 2006). A subset of GS that were re-contacted as part of a mental health follow-up study, are used here for the current investigation (L. B. Navrady et al., 2017). Participants provided information on past 6 month life events and MDD status. We will explore the genetic relationship between MDD, neuroticism and SLE by using GWAS summary statistics from external datasets: the Psychiatric Genetic Consortium (PGC) (MDD) and the Social Science Genetic Association Consortium (SSGAC) (neuroticism).

## Materials and Methods

### Sample Description

The individuals used in this study were a subset of Generation Scotland: the Scottish Family Health Study (GS) which has been described in detail elsewhere (Amador et al., 2015; Nagy et al., 2017; Smith et al., 2013; Smith et al., 2006). Briefly, GS comprises 23,690 individuals aged 18 years and over recruited via general practitioners’ throughout Scotland. In 2014, re-contact of GS participants began as part of a data collection initiative designed to re-assess the mental health of participants. In total, 21,526 GS participants were re-contacted by post and asked to return a questionnaire by post or via a URL link to complete online. Nine thousand six hundred and eighteen participants volunteered as part of the mental health follow-up (45% response rate), and these are the participants used in this study. A full description of the re-contact procedure and data collected is provided elsewhere (L. Navrady et al., 2017). All components of GS, including its protocol and written study materials have received national ethical approval from the NHS Tayside committee on research ethics (reference 05/s1401/89).

SLEs were assessed using the List of Threatening Experiences (Brugha, Bebbington, Tennant, & Hurry, 1985) which is a self-report questionnaire consisting of 12 life events that have taken place in the past six months. In order to perform heritability analyses of SLE we created a list of personal life events that should be unique to the individual endorsing them. This was to prevent a potential inflation in heritability estimates from family members endorsing the same life events (e.g. death of a family member) (Supplementary Table 1). For the remainder of the analyses presented in this study we used total SLE’’s reported and divided the experiences into independent and dependent life events (Brugha et al., 1985) (Supplementary Table 2).

In the GS mental health cohort, lifetime MDD was assessed using the Composite International Diagnostic Interview – Short Form (CIDI-SF) (Kessler, Zhao, Blazer, & Swartz, 1997). The CIDI-SF is a self-report measure of psychiatric symptoms and allows for the ascertainment of lifetime MDD status, age of onset and number of episodes. Neuroticism was assessed during the initial contact of the full GS cohort using the Eysenck personality questionnaire (Eysenck & Sybil, 1975).

Genome-wide genotype data generated using the Illumina Human OmniExpressExome-8-v1.0 array and was available for 8734 of the 9618 individuals from the GS subset. Genotyping is described in greater detail elsewhere (Smith et al., 2006). Population outliers were identified and removed from the sample (Amador et al., 2015). Quality control of genotypes removed SNPs with a call rate < 98%, a missing rate per individual ≥2%, a minor allele frequency (MAF) >1% and Hardy-Weinberg equilibrium (HWE) p ≤ 1×10^−6^. In total, 561,125 autosomal SNPs remained and were used in subsequent analyses. Multidimensional scaling (MDS) components were created according to the ENIGMA 1000 genomes protocol (ENIGMA, 2013) in the software package PLINK (Purcell et al., 2007).

### Heritability analyses

Only personal SLEs (Supplementary Table 1) were used to estimate the heritability of life events. If individuals in a family endorse the same event (e.g. death of a family member) it will not be clear if the similarities between family members are due to endorsement of the same event or shared genetic effects influencing the reporting of SLE. Furthermore, as heritability estimates in family studies can be distorted by shared environments as well as shared genetic material, we estimated heritability whilst modelling components of the environment (Xia et al., 2016). Genetic effects were estimated in GCTA by fitting a pedigree kinship matrix (K) and a genetic relationship matrix (G) alongside 3 environmental components: the environmental effect from the nuclear family F, the environmental effect from the couple relationship C and the environmental effect from the full sibling relationship S. The population prevalence used to transfer heritability estimates for MDD from the observed scale to the liability scale was 0.162 (Kessler et al., 2003; Yang, Lee, Goddard, & Visscher, 2011). The variance explained by these effects were estimated using linear mixed models (LMM) and the statistical significance tested using likelihood ratio (LRT) and Wald tests. Details on the construction of the variance-covariance matrices can be found in the supplemental material.

Genomic and environmental relationship components are fitted in a LMM implemented in GCTA:

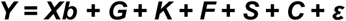

Y is a vector of either a binary MDD phenotype or the score for SLEs. b is the effect of X, a vector of fixed effect covariates which include age, sex and 20 principal components derived from the genome-wide GRM. G and K represent the random genetic effects from the SNPs and the pedigree, respectively. F, S, and C and ε represent the random environmental effects shared by nuclear family members, full-siblings, couples and the error term respectively.

Backward stepwise model selection was used to select the appropriate model to identify major genetic and/or environmental components contributing to the phenotypic variance. The initial model was the full ‘GKFSC’ model and LRT and Wald tests were conducted to test each variance component. A variance component was removed if it failed to obtain significance (α=5%) in both tests and among the variance components satisfying (1) it has the highest P value in the Wald test. This process was repeated until all the remaining components were significant in either the LRT or Wald test. This method is described in more detail by Xia et al (Xia et al., 2016).

There were 659 couple pairs, 1928 full sibling pairs and 4523 nuclear families (containing at least 2 individuals) in the present sample. The number of non-zero elements of the KFSC matrices for whom genotypic and all phenotypic information are available in the present sample are shown in Supplementary Table 3. The G matrix does not contain any non-zero elements.

### Polygenic risk score (PRS) analyses and phenotypic correlations

Polygenic risk scores (PRS) were created by using the raw genotype data from a target sample (GS) and summary statistics from a discovery sample in PRSice software (Euesden, Lewis, & O’Reilly, 2015). This method calculates the sum of associated alleles an individual carries across the genome, weighted by their effect size in an independent GWAS. SNPs were linkage disequilibrium pruned using clump-based pruning (r^2^=0.1, 250□kb window) prior to creating PRS. Scores were created for a range of p-value thresholds ranging from p ≤ 0.01 to p =1 in 0.01 increments. Only one PRS was used to test for association and this was based on which p-value threshold score explained most variance in the trait of interest. The p-value thresholds used are shown in Supplementary Table 4.

PRS were created for MDD (MDD-PRS) and neuroticism (N-PRS). The GWAS summary statistics used for MDD were those from the unpublished Psychiatric Genetics Consortium (PGC MDD29) GWAS of MDD (130,664 cases vs 330,470 controls). For neuroticism PRS, the summary statistics from the Social Science Genetic Association Consortium (SSGAC) GWAS of 170,911 individuals were used (Okbay et al., 2016). Eighteen association tests were carried out between the MDD-PRS/N-PRS and traits of interest, which gave the Bonferroni corrected p-value of 0.0028 as the threshold for statistical significance (tests presented in Table 3 and Table 4).

**Table 3).**
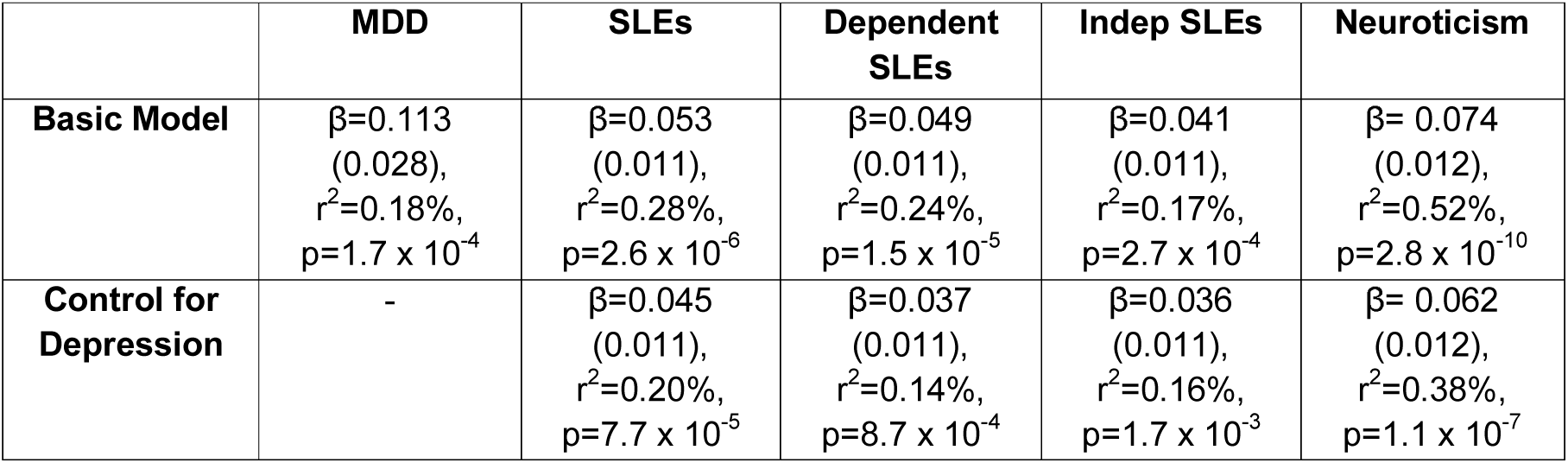
MDD PRS association analyses. Basic model has age, sex and 4 MDS components to control for population stratification and PRS as fixed effects. Family structure was controlled for using a pedigree matrix in AS-Reml. Depression was added as fixed effects in subsequent models. Best threshold PRS for each trait used, for MDD p ≤ 0.52, SLEs p ≤ 0.35 and Neuroticism p ≤ 0.23.

|  | MDD | SLEs | Dependent SLEs | Indep SLEs | Neuroticism |
| --- | --- | --- | --- | --- | --- |
| **Basic Model** | $\beta=0.113$<br>(0.028),<br>$r^2=0.18\%$,<br>$p=1.7 \times 10^{-4}$ | $\beta=0.053$<br>(0.011),<br>$r^2=0.28\%$,<br>$p=2.6 \times 10^{-6}$ | $\beta=0.049$<br>(0.011),<br>$r^2=0.24\%$,<br>$p=1.5 \times 10^{-5}$ | $\beta=0.041$<br>(0.011),<br>$r^2=0.17\%$,<br>$p=2.7 \times 10^{-4}$ | $\beta=0.074$<br>(0.012),<br>$r^2=0.52\%$,<br>$p=2.8 \times 10^{-10}$ |
| **Control for Depression** | - | $\beta=0.045$<br>(0.011),<br>$r^2=0.20\%$,<br>$p=7.7 \times 10^{-5}$ | $\beta=0.037$<br>(0.011),<br>$r^2=0.14\%$,<br>$p=8.7 \times 10^{-4}$ | $\beta=0.036$<br>(0.011),<br>$r^2=0.16\%$,<br>$p=1.7 \times 10^{-3}$ | $\beta=0.062$<br>(0.012),<br>$r^2=0.38\%$,<br>$p=1.1 \times 10^{-7}$ |

**Table 4).**
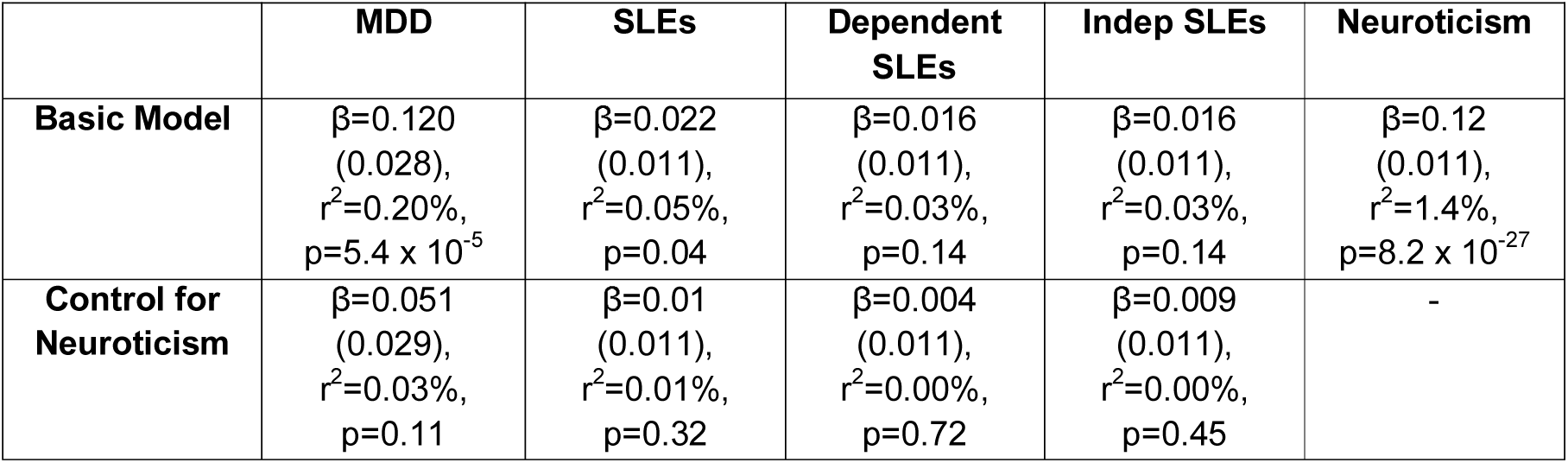
Neuroticism PRS association analyses. Basic model has age, sex and 4 MDS components to control for population stratification and PRS as fixed effects. Family structure was controlled for using a pedigree matrix in AS-Reml. Neuroticism was added as a fixed effect covariate in the subsequent model. Best threshold PRS for each trait used, for MDD p ≤ 0.10 and SLEs/Neuroticism p ≤ 0.60.

|  | **MDD** | **SLEs** | **Dependent SLEs** | **Indep SLEs** | **Neuroticism** |
| --- | --- | --- | --- | --- | --- |
| **Basic Model** | $\beta=0.120$<br>(0.028),<br>$r^2=0.20\%$,<br>$p=5.4 \times 10^{-5}$ | $\beta=0.022$<br>(0.011),<br>$r^2=0.05\%$,<br>$p=0.04$ | $\beta=0.016$<br>(0.011),<br>$r^2=0.03\%$,<br>$p=0.14$ | $\beta=0.016$<br>(0.011),<br>$r^2=0.03\%$,<br>$p=0.14$ | $\beta=0.12$<br>(0.011),<br>$r^2=1.4\%$,<br>$p=8.2 \times 10^{-27}$ |
| **Control for Neuroticism** | $\beta=0.051$<br>(0.029),<br>$r^2=0.03\%$,<br>$p=0.11$ | $\beta=0.01$<br>(0.011),<br>$r^2=0.01\%$,<br>$p=0.32$ | $\beta=0.004$<br>(0.011),<br>$r^2=0.00\%$,<br>$p=0.72$ | $\beta=0.009$<br>(0.011),<br>$r^2=0.00\%$,<br>$p=0.45$ | - |

All variables were log transformed towards normality where necessary. Continuous variables were scaled to have a mean of 0 and a standard deviation of 1, such that the reported regression coefficients (betas) are standardized. Mixed linear models implemented in the ASReml-R (www.vsni.co.uk/software/asreml) software package were used to test the association between MDD-PRS and traits of interest. When associations between binary traits and PRS are reported, Taylor series transformation was used to convert beta and standard error values from the linear scale to the liability scale (International Genetics of Ankylosing Spondylitis et al., 2013). Age, sex and four MDS components were fitted as fixed effect covariates. To control for family structure pedigree information was used to create an inverse relationship matrix which was fitted as a random effect. Wald’s conditional F-test was used to calculate the significance of fixed effects. This method was also used to test the phenotypic association between MDD, SLEs and neuroticism. Relative risk ratios were determined using the R package epitools (https://CRAN.R-project.org/package=epitools).

### LD score regression

To quantify the degree of genetic overlap in common variants between SLEs and PGC-MDD/SSGAC-neuroticism we used LD score regression (Bulik-Sullivan et al., 2015). This method analyses the correlational structure of LD between SNPs and the patterns of association between SNPs and traits of interest to calculate genetic correlations. We performed GWAS of independent and dependent life events in the present GS sample to generate summary statistics for LD score regression. GWAS was performed using mixed linear model association analyses in GCTA using imputed genotype data, implementing a leave-one-chromosome-out approach, which creates a genetic relationship matrix (GRM) excluding the chromosome on which the candidate SNP tested for association is located (Yang, Zaitlen, Goddard, Visscher, & Price, 2014). Fitting a GRM controlled for family structure within the GS sample. Sex, age and 20 MDS components were fitted as fixed effect covariates. Genotypes were imputed using the Haplotype Reference Consortium (HRC) reference panel. Individuals with missingness ≥3% were excluded along with SNPs with a call rate ≤98%, HWE p-value ≤ 1 × 10^−6^ and a MAF ≤ 1%. Genotype SNP data were phased using SHAPEIT2 and imputation performed using PBWT software(Durbin, 2014). Post-imputation SNPs with more than two alleles, monomorphic SNPs and SNPs with an INFO score < 0.8 were removed. QQ plots for the GWAS of independent and dependent life events are shown in the supplementary material.

## Results

The prevalence of MDD in the GS subsample studied here was 16.4% (1506 cases vs 7667 controls). Individuals with a lifetime diagnosis of MDD had significantly higher neuroticism scores, were significantly younger, and were more likely to be female (Table 1). A significant positive association was found between the number of past 6 month stressful life events (SLEs) and MDD (β =0.21, r^2^=1.1%, p=2.5 × 10^−25^); with individuals with MDD reporting, on average, 1.14 SLEs compared to controls who reported an average of 0.83 life events (Table 1). MDD status significantly associated with both dependent (β =0.25, r^2^=1.0%, p=1.8 × 10^−21^) and independent life events (β =0.14, r^2^=0.28%, p=5.3 × 10^−07^). The relative risk (RR) for MDD in individuals experiencing any SLE was 1.44 (95% C.I.-1.31-1.58). The RR risk for MDD peaked in individuals reporting 4 SLEs compared to individuals reporting no life events (RR=1.91, 95% C.I.=1.50-2.44) (Supplementary Figure 1). Neuroticism was significantly and positively associated with SLE (β =0.11, r^2^=1.3%, p=4.60 × 10^−26^), with associations observed for both dependent (β=0.10, r^2^=1.0%, p=2.4 × 10^−21^) and independent (β=0.08, r^2^=0.71%, p=5.1 × 10^−15^) life events.

**Table 1).** Summary of individuals from GS follow up cohort with phenotypic information available. All differences between cases and control significant at p ≤ 5.13 × 10^−12^ after controlling for family structure using a pedigree matrix in AS-Reml.

|  | Cases (N=1506) | Controls (N=7667) |
| --- | --- | --- |
| % Female | 76% | 59.8% |
| Age (s.d.) | 54.2 (12.4) | 56.8 (13.5) |
| SLE Total (s.d.) | 1.14 (1.45) | 0.83 (1.25) |
| Neuroticism | 5.35 (3.46) | 3.15 (2.87) |

To test the heritability of SLE only personal life events were included. These are events that should be unique to an individual. In a family based sample the sum of the G (genetic) and K (pedigree) effects are equivalent to the narrow sense heritability of a trait, when controlling for shared environment (Xia et al., 2016). For personal SLEs, the narrow sense heritability estimate was 0.13 (G=0.07(S.E.=0.04 + K=0.06(S.E.=0.12), but only the SNP genetic effects (G) were statistically significant (p=0.007 and p=0.5 respectively) (Table 2). Using backward stepwise model selection, 8% of the variance in personal SLEs were explained by common genetic effects (S.E.=3%, p=9 × 10^−4^). A significant couple effect was also detected, 13% (S.E.=0.05, p=0.016) (Table 3).

**Table 2).** Partitioning phenotypic variance into environmental and genetic components using the full GKFSC model. Backward stepwise selection was used to select the most parsimonious model for each trait. *Model non-convergence, unconstrained REML performed. Bold values are have significant LRT at p < 0.05.

| Model | G (S.E.) | K (S.E.) | F (S.E.) | C (S.E.) | S (S.E.) |
| --- | --- | --- | --- | --- | --- |
| **Personal SLEs** |  |  |  |  |  |
| GKFCS | **0.07 (0.04)** | 0.06 (0.12) | 0.00 (0.06) | 0.14 (0.08) | 0.00 (0.03) |
| GC | **0.08 (0.03)** |  |  | **0.13 (0.05)** |  |
| **Neuroticism** |  |  |  |  |  |
| GKFCS | **0.11 (0.05)** | 0.10 (0.12) | 0.03 (0.06) | 0.00 (0.07) | 0.01 (0.03) |
| GK | **0.12 (0.05)** | **0.20 (0.06)** |  |  |  |
| **MDD** |  |  |  |  |  |
| GKFCS* | 0.16 (0.10) | -0.23 (0.28) | 0.18 (0.14) | 0.08 (0.18) | 0.05 (0.08) |
| F |  |  | **0.18 (0.04)** |  |  |

We were not able to replicate a previous study by Zeng et al. on the full GS sample (N=19,896), that found shared genetics and couple-associated environment explain 61% of the variance in MDD in the total GS sample (K= 0.35(S.E.=0.06), G= 0.12(S.E.=0.05), C=0.14(S.E.=0.07)) (Zeng et al., 2016). In our sub-sample, the genetic effects on MDD, G and K, were not significant. Our study uses a subset individuals from the Zeng et al. study, and in the present sample only a significant effect of family was detected, but this may be due to reduced power in a sample of only 1506 MDD cases. Using the GCTA power calculator we estimated that we had only 34% power to detect a SNP genetic effect of 0.12 in the GS mental health follow-up cohort (http://cnsgenomics.com/shiny/gctaPower/). The narrow sense heritability estimates for neuroticism was estimated at 0.32, with 12% (S.E.=0.05) of the variance explained by common SNPs (G). The environmental components did not contribute to any of the phenotypic variance in neuroticism and this is in accordance with the findings for neuroticism on the full GS sample reported by Hill et al who found the narrow-sense heritability of neuroticism to be 0.30 (G=0.11 (S.E.=0.02), K= 0.29 (S.E.=0.03) (Hill et al., 2017) (Table 2).

Genetic overlap between SLEs and MDD/neuroticism was tested using PRS. For these analyses we tested the association with total, and also independent and dependent life events. Dependent life events have shown greater association at the phenotypic level with MDD (Harkness & Luther, 2001; Harkness et al., 1999). MDD-PRS were significantly associated with MDD (β=0.11, r2=0.18%, p=1.7 × 10^−4^) and neuroticism (β=0.074, r^2^=0.52%, p=2.8 × 10^−10^) (Table 4). MDD-PRS were also associated with total SLEs (β=0.053 r^2^=0.28%, p=2.6 × 10^−6^). Individuals with a higher MDD PRS report more SLEs. The effect was similar for dependent life events (β=0.049, r^2^=0.24%, p=1.5 × 10^−5^) compared to independent life events (β=0.041, r^2^=0.17%, p=2.7 × 10^−4^) (Table 3). After controlling for MDD status, the association between polygenic risk for MDD and SLEs was still significant although the effect was attenuated (β=0.053 vs β=0.045). This suggests that the association is not driven solely by the increased presence of lifetime MDD in individuals with higher SLE scores. These findings were supported by the results of the LD score regression analyses. There was a significant genetic overlap between total SLEs and MDD (r_G_=0.33, S.E.=0.08) and the genetic correlation was significantly stronger (Z=1.95, p=0.023) for dependent SLEs (r_G_=0.61, S.E.=0.20) compared to independent SLE (r_G_=0.19, S.E.=0.08) (Table 4).

Genetic overlap between SLEs and neuroticism was tested using neuroticism PRS (N-PRS). N-PRS were associated with neuroticism (β=0.12, r^2^=1.4%,p=8.2 × 10^−27^) and MDD (β=0.12, r^2^=0.2%, p=5.4 × 10^−5^). N-PRS were nominally associated with total SLEs (β=0.022, r^2^=0.05%, p=0.04); however, the association was weaker compared to MDD-PRS (β=0.053, r^2^=0.38%, p=2.6 × 10^−6^) and not significant after correction for multiple testing. The association between independent or dependent SLEs and N-PRS were not significant and after controlling for neuroticism the association with SLEs became weaker (Table 5). A significant genetic overlap between neuroticism and total reported SLEs was detected (r_G_=0.15, S.E.=0.07) using LD score regression (Table 5). The genetic correlation between neuroticism and dependent SLEs was 0.25 (S.E.=0.10), but this was not significantly greater (Z=1.56, p=0.06) than the genetic correlation with independent SLEs. The genetic correlation between neuroticism and independent SLEs was not significant.

**Table 5).** Genetic correlation (rG) between SLEs and MDD using LD score regression (Bulik-Sullivan et al, 2015). All estimates in bold are statistically significant p ≤ 0.05. PGC-MDD GWAS summary statistics taken from PGC GWAS of MDD (MDD29).

|  | **PGC-MDD (S.E.)** | **Neuroticism (S.E.)** |
| --- | --- | --- |
| **Total SLEs** | **0.33 (0.08)** | **0.15 (0.07)** |
| **Independent SLEs** | **0.19 (0.08)** | 0.06 (0.07) |
| **Dependent SLEs** | **0.61 (0.20)** | **0.25 (0.10)** |

## Discussion

Using a polygenic risk score (PRS) approach MDD and SLEs were found to have shared polygenic architecture. MDD polygenic risk was found to be higher in individuals reporting more SLEs. LD score regression showed a genetic correlation between MDD and SLEs using summary statistics from an independent MDD cohort. We also report a positive genetic correlation between neuroticism and SLEs. The variance in reporting of personal SLEs can be partly explained by common SNP effects and the environment shared by couples. 8% of the variance in personal SLEs was attributable to common genetic variants and an additional 13% of was explained by couple shared environment. This left 79% of the variance in personal SLEs unexplained by common genetic or shared environmental effects.

The narrow sense heritability point estimate for personal SLEs in the current sample was 13% which is lower than the 20-50% range of estimates derived from twin studies (Billig et al., 1996; Foley et al., 1996; Kendler et al., 1993; Plomin et al., 1990). Furthermore, the pedigree contribution to this effect was not statistically significant. When personal SLEs were analysed modelling both genetic and environmental components, the SNP heritability estimate was significant and accounted for 8% of the variance in SLEs. This is the same as the estimate derived from the population-based study of African American women that found the SNP heritability of SLEs to be 8% (Dunn et al., 2016). However, another study found SNP effects account for roughly a third of variance in SLEs (Power et al., 2013). This is in contrast to our own findings and those of Dunn et al. and may be due to the high proportion of clinically ascertained MDD cases in the Power et al. sample (Dunn et al., 2016; Power et al., 2013). As MDD and SLEs are genetically correlated this may inflate heritability estimates if samples have a high proportion of MDD cases. In the present study we model genetic and environmental influences using different types of relationships and find that the heritability of SLEs are much lower than is often reported in twin studies.

We also detected a significant effect of the environment shared by couples on personal SLEs. The effect of couple shared environment on variance in MDD has previously been reported on the full GS cohort (Zeng et al., 2016) to be 15-22%. We find that 13% of the variance in self-reported SLEs in this sample is attributable to shared couple environment. A study of 354 male Vietnam era veterans found that spousal correlations in depression were due to common stressors and that there were crossover effects so that depression in one spouse was influenced by stressors reported by the other (Westman & Vinokur, 1998). Our data support this finding and reinforces the importance of recent shared environment on MDD and SLEs. We find little evidence for the effect of nuclear family or sibling environment on reporting SLEs. A recent study of anthropometric and cardiometabolic traits in GS found that ~11% of variation across traits could be explained by the environment common to couples (Xia et al., 2016) suggesting that recent shared environment is important when modelling the heritability of complex diseases. However, it should be noted that there might be assortative mating between spouses in which case modelling the couple correlation entirely as an environmental effect may inflate heritability estimates (Vinkhuyzen, van der Sluis, Maes, & Posthuma, 2012).

A significant genetic correlation between SLEs and MDD was identified in this sample. PRS for MDD were associated with both dependent and independent SLEs even after controlling for MDD status. Using LD score regression, we found that the genetic overlap between dependent life events and MDD (0.61) was higher than for independent life events (0.19). This is in line with the findings by Dunn et al. who found a strong genetic correlation between MDD and SLEs in a sample of African American women (r_G_=0.95) (Dunn et al., 2016). The genetic overlap between SLEs and MDD calls for a different interpretation of the effect of SLEs on MDD. Rather than considering SLEs simply as risk factors for MDD, the SNPs which predispose to MDD also increase risk for SLEs. This may arise from individuals selecting themselves into high risk stress environments or via personality traits, such as neuroticism, which prime them to respond negatively to life events (Kendler et al., 2004) or higher reporting. We found a significant genetic correlation between neuroticism and SLEs that was more pronounced for dependent SLEs but not independent SLEs. This supports previous studies which have shown that neuroticism is associated with increased reporting and sensitivity to SLEs (Gautam & Kamal, 1990; Kendler et al., 2004; Magnus et al., 1993). The discrepancy between the N-PRS and the LD score regression analyses is likely due to the small amount of variance that can be explained by PRS.

There are a number of limitations to this study. Firstly, we rely on self-reported measures of MDD and SLEs which are subject to recall bias. However, a recent study of GS found self-reported and SCID defined MDD to be highly genetically correlated (Zeng et al., 2016) although the bias in the reporting of SLE is unknown. Secondly, the full GKSFC model has its own limitations as a number of the matrices will be correlated such as the nuclear family matrix and the sibling matrix. This could prevent accurate estimates of familial effects. In order to account for this, we performed backward stepwise selection to select the most influential components to each trait however a superior approach would be to use a much larger sample size with more familial relationships. In our case, we were limited by the number of participants in our follow-up mental health study, and the familial structure within this sub-sample of GS. We did not have power to detect common SNP genetic effects for MDD in this sample. Our study suggests that the SNP heritability of personal SLEs are likely to be low and therefore larger samples are warranted to investigate this further. Finally, determining the familial environmental effects on SLEs is challenging when families will endorse the same events solely because they have occurred within the same social network, such as ‘did a close relative of yours die?’. This is also true for couples where major financial crises will be reported by both spouses due to shared assets. We attempted to control for this by creating a personal SLE category and also excluding events that could be inferred by spouses, however people may still endorse an event that happens to a spouse or family member as their own as they find it to be stressful to themselves. It is not possible to ascertain with the data available from this cohort, whether events endorsed by members of a couple reflect the same event, or whether each individual experiences an independent event.

In conclusion, we provide evidence that personal SLEs are heritable but that the effect attributable to common genetic SNPs is likely to be small. The recent environment such as that shared by couples is also likely to contribute to SLEs. There is strong genetic overlap between MDD and SLEs and some genetic overlap between neuroticism and SLEs. These findings underlie the importance of appropriately modelling environmental effects when studying these traits. Furthermore, our results demonstrate that the relationship between SLEs, MDD and personality may not be directionally causal, but a consequence of common genetic effects that influence these traits.

## Acknowledgements

We are grateful to the families who took part in GS, the GPs and Scottish School of Primary Care for their help in recruiting them, and the whole GS team, which includes academic researchers, clinic staff, laboratory technicians, clerical workers, IT staff, statisticians and research managers. Generation Scotland received core support from the Chief Scientist Office of the Scottish Government Health Directorates [CZD/16/6] and the Scottish Funding Council [HR03006]. Genotyping of the GS samples was carried out by the Genetics Core Laboratory at the Wellcome Trust Clinical Research Facility, Edinburgh, Scotland and was funded by the Medical Research Council UK and the Wellcome Trust (Wellcome Trust Strategic Award “STratifying Resilience and Depression Longitudinally” (STRADL) Reference 104036/Z/14/Z). We acknowledge with gratitude the financial support received for this work from the Dr Mortimer and Theresa Sackler Foundation. PT, DJP, IJD, and AMM are members of The University of Edinburgh Centre for Cognitive Ageing and Cognitive Epidemiology, part of the cross-council Lifelong Health and Wellbeing Initiative (MR/K026992/1). Funding from the Biotechnology and Biological Sciences Research Council and Medical Research Council is gratefully acknowledged. CSH, CA, CX and PN acknowledge funding from the MRC UK (grants MC_PC_U127592696 and MC_PC_U127561128).

## References

Amador, C., Huffman, J., Trochet, H., Campbell, A., Porteous, D., Wilson, J. F., … Haley, C. S. (2015). Recent genomic heritage in Scotland. BMC Genomics, 16, 437. doi:10.1186/s12864-015-1605-2 10.1186/s12864-015-1605-2 [pii]

Bemmels, H. R., Burt, S. A., Legrand, L. N., Iacono, W. G., & McGue, M. (2008). The heritability of life events: an adolescent twin and adoption study. Twin Res Hum Genet, 11(3), 257–265. doi:10.1375/twin.11.3.257 10.1375/twin.11.3.257 [pii]

Billig, J. P., Hershberger, S. L., Iacono, W. G., & McGue, M. (1996). Life events and personality in late adolescence: genetic and environmental relations. Behav Genet, 26(6), 543–554.

Boardman, J. D., Alexander, K. B., & Stallings, M. C. (2011). Stressful life events and depression among adolescent twin pairs. Biodemography Soc Biol, 57(1), 53–66.

Brugha, T., Bebbington, P., Tennant, C., & Hurry, J. (1985). The List of Threatening Experiences: a subset of 12 life event categories with considerable long-term contextual threat. Psychol Med, 15(1), 189–194.

Bulik-Sullivan, B. K., Loh, P. R., Finucane, H. K., Ripke, S., Yang, J., Patterson, N., … Neale, B. M. (2015). LD Score regression distinguishes confounding from polygenicity in genome-wide association studies. Nat Genet, 47(3), 291–295. doi:10.1038/ng.3211 [pii] 10.1038/ng.3211

Chun, C. A., Cronkite, R. C., & Moos, R. H. (2004). Stress generation in depressed patients and community controls. Journal of Social and Clinical Psychology., 23, 392–412.

Dunn, E. C., Wiste, A., Radmanesh, F., Almli, L. M., Gogarten, S. M., Sofer, T., … Smoller, J. W. (2016). Genome-Wide Association Study (Gwas) and Genome-Wide by Environment Interaction Study (Gweis) of Depressive Symptoms in African American and Hispanic/Latina Women. Depress Anxiety, 33(4), 265–280. doi:10.1002/da.22484

Durbin, R. (2014). Efficient haplotype matching and storage using the positional Burrows-Wheeler transform (PBWT). Bioinformatics, 30(9), 1266–1272. doi:10.1093/bioinformatics/btu014

Enigma, T. E. N. G. t. M. A. E. C. (2013). ENIGMA2 Genetics Support Team ENIGMA2 1KGP Cookbook (v3). [Online].

Euesden, J., Lewis, C. M., & O’Reilly, P. F. (2015). PRSice: Polygenic Risk Score software. Bioinformatics, 31(9), 1466–1468. doi:10.1093/bioinformatics/btu848 [pii] 10.1093/bioinformatics/btu848

Eysenck, H. J., & Sybil, B. G. (1975). Manual of the Eysenck Personality Questionnaire. London: Hodder and Stoughton.

Foley, D. L., Neale, M. C., & Kendler, K. S. (1996). A longitudinal study of stressful life events assessed at interview with an epidemiological sample of adult twins: the basis of individual variation in event exposure. Psychol Med, 26(6), 1239–1252.

Gautam, S., & Kamal, P. (1990). A study of impact of stressful life-events in neurotic patients. Indian J Psychiatry, 32(4), 356–361.

Glahn, D. C., Curran, J. E., Winkler, A. M., Carless, M. A., Kent, J. W., Jr., Charlesworth, J. C., … Blangero, J. (2012). High dimensional endophenotype ranking in the search for major depression risk genes. Biol Psychiatry, 71(1), 6–14. doi:10.1016/j.biopsych.2011.08.022

Hammen, C. (1991). Generation of stress in the course of unipolar depression. J Abnorm Psychol, 100(4), 555–561.

Harkness, K. L., & Luther, J. (2001). Clinical risk factors for the generation of life events in major depression. J Abnorm Psychol, 110(4), 564–572.

Harkness, K. L., Monroe, S. M., Simons, A. D., & Thase, M. (1999). The generation of life events in recurrent and non-recurrent depression. Psychol Med, 29(1), 135–144.

Hill, W. D., Arslan, R., Xia, C., Luciano, M., Amador, C., Navarro, P., … Penke, L. (2017). Genomic analysis of family data reveals additional genetic effects on intelligence and personality. (http://biorxiv.org/content/early/2017/02/06/106203).

International Genetics of Ankylosing Spondylitis, C., Cortes, A., Hadler, J., Pointon, J. P., Robinson, P. C., Karaderi, T., … Brown, M. A. (2013). Identification of multiple risk variants for ankylosing spondylitis through high-density genotyping of immune-related loci. Nat Genet, 45(7), 730–738. doi:10.1038/ng.2667

Jardine, R., Martin, N. G., & Henderson, A. S. (1984). Genetic covariation between neuroticism and the symptoms of anxiety and depression. Genet Epidemiol, 1(2), 89–107. doi:10.1002/gepi.1370010202

Jylha, P., & Isometsa, E. (2006). The relationship of neuroticism and extraversion to symptoms of anxiety and depression in the general population. Depress Anxiety, 23(5), 281–289. doi:10.1002/da.20167

Kendler, K. S., & Gardner, C. O. (2010). Dependent stressful life events and prior depressive episodes in the prediction of major depression: the problem of causal inference in psychiatric epidemiology. Arch Gen Psychiatry, 67(11), 1120–1127. doi:67/11/1120 [pii] 10.1001/archgenpsychiatry.2010.136

Kendler, K. S., & Karkowski-Shuman, L. (1997). Stressful life events and genetic liability to major depression: genetic control of exposure to the environment? Psychol Med, 27(3), 539–547.

Kendler, K. S., Karkowski, L. M., & Prescott, C. A. (1999). Causal relationship between stressful life events and the onset of major depression. Am J Psychiatry, 156(6), 837–841. doi:10.1176/ajp.156.6.837

Kendler, K. S., Kessler, R. C., Walters, E. E., MacLean, C., Neale, M. C., Heath, A. C., & Eaves, L. J. (1995). Stressful life events, genetic liability, and onset of an episode of major depression in women. Am J Psychiatry, 152(6), 833–842. doi:10.1176/ajp.152.6.833

Kendler, K. S., Kuhn, J., & Prescott, C. A. (2004). The interrelationship of neuroticism, sex, and stressful life events in the prediction of episodes of major depression. Am J Psychiatry, 161(4), 631–636. doi:10.1176/appi.ajp.161.4.631

Kendler, K. S., Neale, M., Kessler, R., Heath, A., & Eaves, L. (1993). A twin study of recent life events and difficulties. Arch Gen Psychiatry, 50(10), 789–796.

Kessler, R. C. (1997). The effects of stressful life events on depression. Annu Rev Psychol, 48, 191–214. doi:10.1146/annurev.psych.48.1.191

Kessler, R. C., Berglund, P., Demler, O., Jin, R., Koretz, D., Merikangas, K. R., … Wang, P. S. (2003). The epidemiology of major depressive disorder: results from the National Comorbidity Survey Replication (NCS-R). JAMA, 289(23), 3095–3105. doi:10.1001/jama.289.23.3095289/23/3095 [pii]

Kessler, R. C., Zhao, S., Blazer, D. G., & Swartz, M. (1997). Prevalence, correlates, and course of minor depression and major depression in the National Comorbidity Survey. J Affect Disord, 45(1–2), 19–30. doi:S0165-0327(97)00056-6 [pii]

Magnus, K., Diener, E., Fujita, F., & Pavot, W. (1993). Extraversion and neuroticism as predictors of objective life events: a longitudinal analysis. J Pers Soc Psychol, 65(5), 1046–1053.

Nagy, R., Boutin, T. S., Marten, J., Huffman, J. E., Kerr, S. M., Campbell, A., … Hayward, C. (2017). Exploration of haplotype research consortium imputation for genome-wide association studies in 20,032 Generation Scotland participants. Genome Med, 9(1), 23. doi:10.1186/s13073-017-0414-4

Navrady, L., Wolters, M. K., Macintyre, D. J., Clarke, T. K., Campbell, A., Murray, A. D., … McIntosh, A. M. (2017). Cohort Profile: Stratifying Resilience and Depression Longitudinally (STRADL): A questionnaire follow-up of Generation Scotland: Scottish Family Health Study (GS:SFHS). International Journal of Epidemiology.

Navrady, L. B., Wolters, M. K., Macintyre, D., Clarke, T.-K., Campbell, A., Murray, A., … McIntosh, A. M. (2017). Cohort Profile: Stratifying Resilience and Depression Longitudinally (STRADL): A questionnaire follow-up of Generation Scotland: Scottish Family Health Study (GS:SFHS). International Journal of Epidemiology, In press.

Okbay, A., Baselmans, B. M., De Neve, J. E., Turley, P., Nivard, M. G., Fontana, M. A., … Cesarini, D. (2016). Genetic variants associated with subjective well-being, depressive symptoms, and neuroticism identified through genome-wide analyses. Nat Genet, 48(6), 624–633. doi:10.1038/ng.3552

Plomin, R., Lichtenstein, P., Pedersen, N. L., McClearn, G. E., & Nesselroade, J. R. (1990). Genetic influence on life events during the last half of the life span. Psychol Aging, 5(1), 25–30.

Power, R. A., Wingenbach, T., Cohen-Woods, S., Uher, R., Ng, M. Y., Butler, A. W., … McGuffin, P. (2013). Estimating the heritability of reporting stressful life events captured by common genetic variants. Psychol Med, 43(9), 1965–1971. doi:10.1017/S0033291712002589 [pii] 10.1017/S0033291712002589

Purcell, S., Neale, B., Todd-Brown, K., Thomas, L., Ferreira, M. A., Bender, D., … Sham, P. C. (2007). PLINK: a tool set for whole-genome association and population-based linkage analyses. Am J Hum Genet, 81(3), 559–575. doi:10.1086/519795

Smith, B. H., Campbell, A., Linksted, P., Fitzpatrick, B., Jackson, C., Kerr, S. M., … Morris, A. D. (2013). Cohort Profile: Generation Scotland: Scottish Family Health Study (GS:SFHS). The study, its participants and their potential for genetic research on health and illness. Int J Epidemiol, 42(3), 689–700. doi:10.1093/ije/dys084 [pii] 10.1093/ije/dys084

Smith, B. H., Campbell, H., Blackwood, D., Connell, J., Connor, M., Deary, I. J., … Morris, A. D. (2006). Generation Scotland: the Scottish Family Health Study; a new resource for researching genes and heritability. BMC Med Genet, 7, 74. doi:10.1186/1471-2350-7-74 [pii] 10.1186/1471-2350-7-74

Surtees, P. G., Miller, P. M., Ingham, J. G., Kreitman, N. B., Rennie, D., & Sashidharan, S. P. (1986). Life events and the onset of affective disorder. A longitudinal general population study. J Affect Disord, 10(1), 37–50. doi: 0165-0327(86)90047-9 [pii]

Vinkhuyzen, A. A., van der Sluis, S., Maes, H. H., & Posthuma, D. (2012). Reconsidering the heritability of intelligence in adulthood: taking assortative mating and cultural transmission into account. Behav Genet, 42(2), 187–198. doi:10.1007/s10519-011-9507-9

Westman, M., & Vinokur, A. D. (1998). Unraveling the Relationship of Distress Levels Within Couples: Common Stressors, Empathic Reactions, or Crossover via Social Interaction? Human Relations, 51(2), 137–156.

Xia, C., Amador, C., Huffman, J., Trochet, H., Campbell, A., Porteous, D., … Haley, C. S. (2016). Pedigree-and SNP-Associated Genetics and Recent Environment are the Major Contributors to Anthropometric and Cardiometabolic Trait Variation. PLoS Genet, 12(2), e1005804. doi:10.1371/journal.pgen.1005804 PGENETICS-D-15-01369 [pii]

Yang, J., Lee, S. H., Goddard, M. E., & Visscher, P. M. (2011). GCTA: a tool for genome-wide complex trait analysis. Am J Hum Genet, 88(1), 76–82. doi:S0002-9297(10)00598-7 [pii] 10.1016/j.ajhg.2010.11.011

Yang, J., Zaitlen, N. A., Goddard, M. E., Visscher, P. M., & Price, A. L. (2014). Advantages and pitfalls in the application of mixed-model association methods. Nat Genet, 46(2), 100–106. doi:10.1038/ng.2876 [pii] 10.1038/ng.2876

Zeng, Y., Xia, C., Amador, C., Fernandez-Pujals, A. M., Thomson, P. A., Campbell, A., … McIntosh, A. M. (2016). Shared genetics and couple-associated environment are major contributors to the risk of both clinical and self-declared depression. EBioMedicine, 14, 161–167.

